# Heparanase-2 protects from endothelial injury by inhibiting TLR4 signaling

**DOI:** 10.1101/705939

**Authors:** Yulia Kiyan, Sergey Tkachuk, Kestutis Kurselis, Nelli Shushakova, Klaus Stahl, Damilola Dawodu, Roman Kiyan, Boris Chichkov, Hermann Haller

**Author notes:** **Contact:** Dr. Yulia Kiyan, Clinic for Nephrology, Hannover Medical School, Carl Neuberg-Strasse 1, 30625 Hannover.

## Abstract

**Objective:** The endothelial glycocalyx and the regulation of its shedding are important to vascular health. Endo-β-D-glucuronidase heparanase-1 (HPSE1) is the only enzyme that can shed heparan sulfate. However, the mechanisms are not well understood.

**Approach and results:** To investigate HPSE1 and its endogenous inhibitor, heparanase-2 (HPSE2), we used cell culture, lentiviral protein overexpression, a microfluidic chip model of cell culture under shear stress conditions, and lipopolysaccharide (LPS) injections in mice. We show that HPSE1 activity aggravated Toll-like receptor 4 (TLR4)-mediated response of endothelial cells to LPS. On the contrary, HPSE2 overexpression was protective. The microfluidic chip flow model confirmed that HPSE2 prevented heparan sulfate shedding by HPSE1. Furthermore, heparan sulfate did not interfere with cluster of differentiation-14 (CD14)-dependent LPS binding, but instead reduced the presentation of the LPS-CD14 complex to TLR4. HPSE2 reduced LPS-mediated TLR4 activation by LPS, subsequent cell signaling, and cytokine expression. Moreover, HPSE2-overexpressing endothelial cells remained protected against LPS-mediated loss of cell-cell contacts. In vivo, expression of HPSE2 in plasma and kidney medullary capillaries was decreased in mouse sepsis model. We next applied purified HPSE2 in mice and observed decreases in TNFα and IL-6 plasma concentrations after intravenous LPS injections.

**Conclusions:** Our data demonstrate the important role of heparan sulfate and the glycocalyx in endothelial cell activation and suggest a protective role of HPSE2 in microvascular inflammation. HPSE2 offers new options for protection against HPSE1-mediated endothelial damage and preventing microvascular disease.

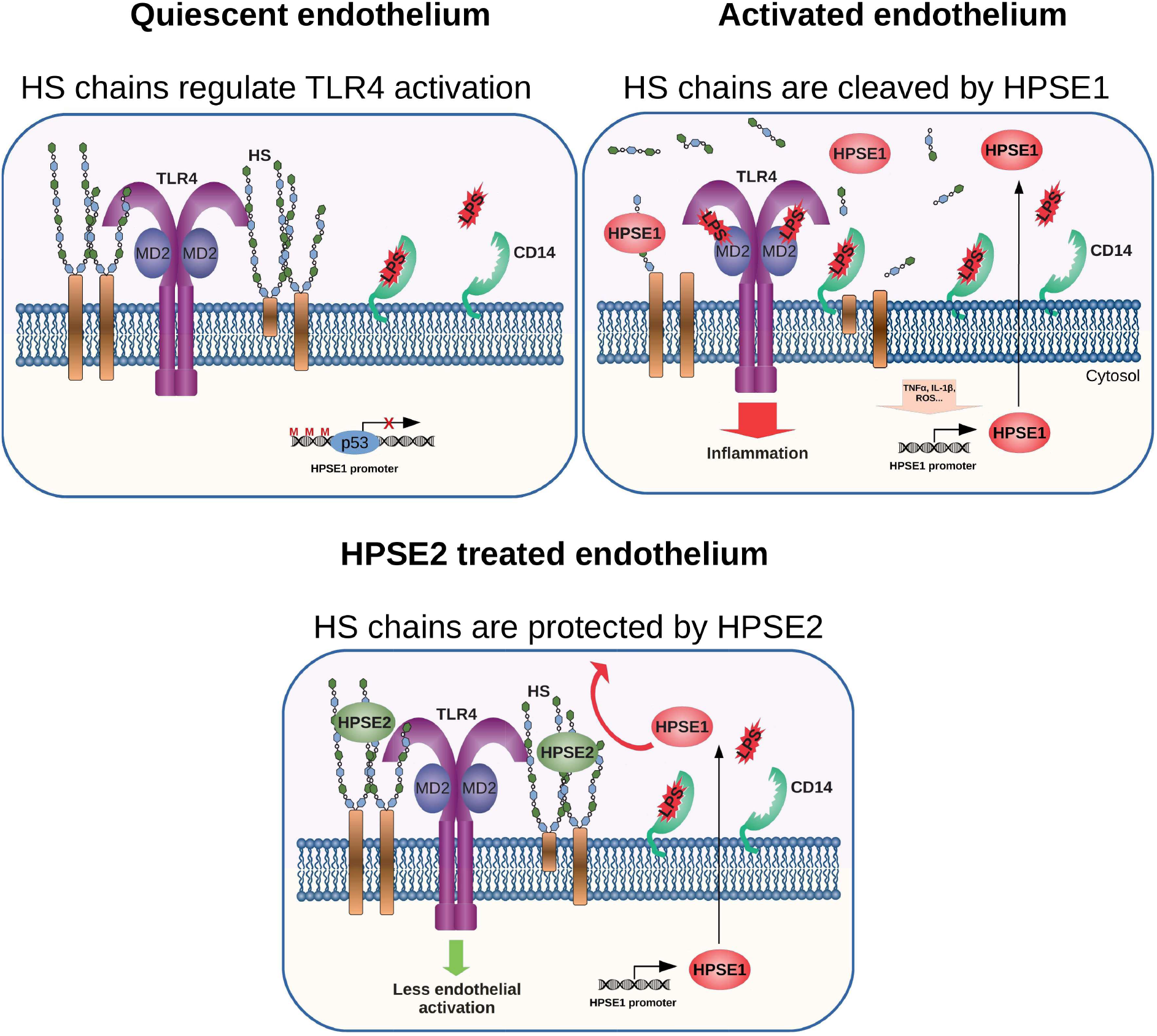
Graphical abstract.

Non standard abbreviations.

3 Non standard abbreviations
|  |  |
| --- | --- |
| HPSE1 | Heparanase 1 |
| HPSE2 | Heparanase 2 |
| LPS | Lipopolysaccharide |
| TLR4 | Toll-like receptor 4 |
| CD14 | Cluster of differentiation 14 |
| PDMS | Polydimethylsiloxan |
| CLP | Cecal ligation and puncture |

## Introduction

The endothelial glycocalyx comprises a network of core proteins and branching glycosaminoglycan polymers such as heparan sulfate, chondroitin sulfate, and hyaluronans, that cover the luminal side of endothelial cells. The glycocalyx is essential to maintain integrity and function of the endothelial layer^1–3^. It serves as a receptor or co-receptor for various ligands such as growth factors, cytokines and chemokines, coagulation factors, proteases, complement-system proteins, membrane receptors, and adhesion proteins^4^. Endothelial glycocalyx shedding occurs during many pathological conditions^5^, including cardiovascular, renal diseases^6,7^ and sepsis^8–10^. Heparan sulfate is a highly sulfated polysaccharide element of the endothelial glycocalyx. Experiments in gene-deleted mice showed that glycocalyx disturbances are not compatible with life.^11,12^. The heparan sulfate interactome is formed by a 300-protein network^4^. Thus, degradation of heparan sulfate affects numerous cellular functions. The only known endo-β-D-glucuronidase capable of degrading heparan sulfate chains in mammals is heparanase 1 (HPSE1). HPSE1 is upregulated in many inflammatory diseases including cancer^13^, diabetes^14^, and sepsis^10^.

The Toll-like receptor-4 (TLR4) is an important inflammatory mediator^15^. Several reports addressed the role of heparan sulfate in the TLR4-mediated response. Soluble heparan sulfate chains serve as TLR4 ligands^16,17^. Cleavage of heparan sulfate by HPSE1 strongly promoted TLR4 response in macrophages^18^. Furthermore, TLR4 signaling was inhibited in cells cultured on heparan sulfate proteoglycans-rich matrix. This inhibition was relieved by the matrix degradation^19^. However in microglia, overexpression of HPSE1 led to diminished LPS-induced response^20^. Co-localization of TLR4 co-receptor Cluster of Differentiation 14 (CD14) with heparan sulfate has been observed that let the authors to suggest that heparan sulfate promotes TLR4/CD14 association. In addition, it has been demonstrated that HPSE1 can act via TLRs causing pro-inflammatory response that is independent of catalytic activity^16,21^.

McKenzie and colleagues cloned and characterized the Heparanase 2 (HPSE2) protein, that exhibits a 40% homology to HPSE1^22^. HPSE2 has no catalytic activity, but instead can inhibit HPSE1^23^. An antagonistic interaction of the heparanases in cancer have been recently reviewed recently^24^. Although HPSE2 circulates in the human plasma^13^, interactions between HPSE1 and HPSE2 in endothelial cells has not been investigated. Endothelial cells express HPSE1 in vitro^25,26^ and serve as an important local source of HPSE1during inflammation^27^. We studied HPSE1 and HPSE2 in endothelial cells and animal models to elucidate the role of these enzymes in maintaining endothelial integrity

## Materials and methods

### 1. Cells, antibody, Real Time-PCR

Human microvascular endothelial cells line HMEC-1 was from ATCC. Cells were cultured as recommended by the supplier. Primary human dermal microvascular endothelial cells were from Promocell. Cells were cultivated as recommended by the supplier and used in the passage 4. Unless otherwise indicated, HMEC-1 cells were used through the study. The following antibodies were used: Heparan sulfate (clone 10e4) from Amsbio; HPSE1 (GTX32650) from GeneTex; Heparanase 2 (NBP1-31457) from Novus Biologicals were used for western blotting; 1c7 monoclonal blocking antibody to HPSE2 was a kind gift of Prof. I. Vlodavsky; antibodies to HPSE2 purchased from Sigma (Cat. # HPA044603) were used for tissue staining; Tubulin (556321) from BD Pharmingen, P-p65 (93H1); P-p38 (D3F9), P-MEK (166M8), and cleaved Caspase 3 antibody (9665) were from Cell Signaling Technologies; VE-cadherin antibody (Clone 123413) from R&D Systems; mouse CD31 Dianova (Dia-310) were used for tissue staining; phalloidin (Alexa 488) was from Invitrogen. DraQ5 was from BioStatus Ltd. OGT 2115 was from Tocris. Purified heparan sulfate (10-70 kDa) was from Sigma. Bovine fibronectin was from Sigma. Purified catalytically active HPSE1 was from R&D Systems.

Cells proliferation was assessed by BrdU Cell proliferation ELISA from Roche. IL-6 ELISA was from Affimetrics. Western blotting was performed as established earlier^28^. Dot blot was performed using nitrocellulose membrane and Heparan sulfate (clone 10e4).

RNA was isolated from cells using Quiagen RNAeasy kits. RT-PCR Taqman assay was performed using Roche TaqMan Master Mix and Roche LightCycler96. The sequence of the primers is given in the Supplementary Table 1. TaqMan Gene Expression Assays for RANTES, IFNB1, and actin as a housekeeper were from Thermofisher Scientific. Human Common Cytokine RT^2^ profiler array (Quiagen) was used accordingly to manufacturer’s instructions. TMB substrate kit was from Thermofisher Scientific.

### 3. Lentivirus for HPSE2 and HPSE1 overexpression

Construction of the expression vectors was performed according to the standard cloning protocols. The vectors were generated by recombination of pDEST12.2 (Invitrogen) with vectors containing full length HPSE2 and HPSE1 (pENRT223.1_HPSE1, pENRT223.1_HPSE2, MyBioSourse, San Diego, CA, USA). Gateway LR Clonase enzyme mix (Invitrogen) catalysis the in vitro recombination between an entry clone pENTR223.1_HPSE2 and destination vector pDEST12.2 to generate expression clones. Correct clones was selected by PstI (New England Biolabs) digestion and confirmed by sequencing (SeqLab). For all cloning experiments in current work subcloning efficiency DH5 alpha competent cells (Invitrogen) were used accordingly to the standard protocols recommended by the supplier.

### 4. Lentiviral gene transfer of HPSE2 and HPSE1

Briefly, the lentivirus packaging genome pCMV-dR8.74^29^, and pMG2G vectors were kindly provided by Dr. Didier Trono (Department of Genetics and Microbiology, Faculty of Medicine, University of Geneva, Switzerland). The lentivirus transfer vector used in this study was generated by Gateway cloning strategy (Invitrogen). An entry vector pENRT223.1 HPSE2 (MyBioSourse) was used. As Gateway Destination vector was used modified plasmid pWPTS-GFP (Tronolab). Modification plasmid pWPTS-GFP was performed by Gateway Vector Conversion System (Invitrogen). A Gateway cassette containing attR recombination sites flanking ccdB gene and a chloramphenicol-resistance gene are blunt-end cloned into cloning site pWPTS-GFP to generate a pWPTS-Dest vector. Gateway LR Clonase enzyme mix (Invitrogen) carried over the in vitro recombination between an entry clone pENTR-HPSE2 and destination vector pWPTS-Dest to generate expression clones pWPTS-HPSE2. pCMV-dR8.74, pMD2G and pWPTS-HPSE2 plasmids were purified using an EndoFree^®^ Plasmid Maxi kit (Qiagen, Valencia, CA), and co-transfected (using ratio pWPTS-HPSE2:pCMV-dR8.74:pMD2G = 3:2:1) into 293T cells using PerFectin transfection reagent (Genlantis) as per manufacturer. After 48 h post transfection cell supernatants, containing viral particles, were filtered using 0.4 mm Steriflip vacuum filtration system (Millipore) and concentrated by ultracentrifugation at 50 000 g for 1.5h at 4°C^30^. Viral titer was determined by LV Lentiviral Titer kit (MoBiTec) and viruses where used at a Multiplicity of infection of 1 – 5 using polybrene (H9268, Sigma) at a concentration of 2 μg/ml.

### 5. Microfluidic experiments

Microfluidic chips were fabricated of polydimethylsiloxan (PDMS) (Sylgard 184, Dow Corning) by replication of the polymeric master. The master was produced at the Institute for Quantum Optics, Leibniz University (Hannover) from diurethane dimethacrylate resin (Sigma-Aldrich) by two-photon polymerization (2PP) technology^31^ The chips consisted of 4 parallel 10mm-long and 2mm-wide channels. The height of the channels was 1 mm. PDMS-microfluidic chips were sealed with 150 μm thick microscopy cover glass using plasma bonding^32^. The diameter of the chip was matched to 35 mm diameter of the cover glass for compatibility with the Okolab incubation chamber.

For operating the microfluidic chips and carrying out cell culture experiments a microfluidic actuation platform was built on the basis of Okolab BOLD LINE H301BL stage incubation set, 4-channel 400VDL/VM4 OEM peristaltic pumps (Watson-Marlow GmbH). Equipment integrated into system provided incubation of the Chips, liquids circulation through the chip, loading/extraction of the cultured cells and loading specific chemicals for experiments. The operation of the microfluidic actuation platform was computer-controlled. The microfluidic interface was based on standard Teflon^®^ tubes with outside diameter of 1.8mm and standard Omni-Lok^®^ microfluidic connectors (Diba Industries Inc.).

Internal surfaces of the microfluidic chip channels were coated with 2μg/ml bovine fibronectin to facilitate cell adhesion. Control virus-transduced cells were seeded in 2 channels of the chip; whereas the other two channels were seeded with HPSE2-overexpressing HMEC-1 cells. Thus, identical experimental conditions were provided for both cell variants. Cells were allowed to adhere for 2-3 hrs, after that the medium flow was initiated in all four channels. Medium flow rate was used form 0.4 ml/h that corresponded to 0.1 dyn/cm^2^ shear stress to 159 ml/h that corresponded to 38.75 dyn/cm^2^. Cells were incubated under flow conditions for 3 days before the experiments. Purified active HPSE1 was perfused through the channels for 4 hrs. Cells were fixed by perfusion with 2% PFA and then stained.

Confocal microscopy was performed using Leica TCS-SP2 AOBS confocal microscope (Leica Microsystems). All the images were taken with oil-immersed x63 objective, NA 1.4. Series of z-scans were processed and quantified using ImageJ software.

### 6. Biotin-LPS binding and pull-down assay

Ultrapure biotin-LPS that activates only TLR4 pathway was purchased from InVivoGen. To analyze LPS binding, cells were placed on ice to prevent receptor internalization and incubated with 50ng/ml biotin-LPS for 1 h. Then cells were washed three times with 1% BSA in PBS and incubated with streptavidin-conjugated HRP on ice for 1 h. HRP activity was quantified using TMB substrate kit. Measurements were take using Tecan plate reader.

For pull down assay strepavidin-magnetic beads were used. Cells were incubated on ice with 50 ng/ml biotin-LPS. Then, the cells were placed at 37°C for indicated time to allow for activation of TLR4 signaling complex. Then cells were placed on ice, washed, and cell lysis was performed. Cell lysates were incubated with streptavidin-beads for 1h at 4°C on rotator, then washed three times with ice cold PBS and used for electrophoresis followed by immunoblotting.

### 7. Cell surface TLR4 ELISA

Cell surface TLR4 expression was measured on live cells incubated on ice in buffer containing 0.5% BSA/PBS with 0.05% NaN3 in ELISA plate^33^. Staining was quantified using HRP-antibody and TMB substrate kit.

### 8. Immunocytochemistry

After the stipulated time, cells grown on coverslips were fixed and processed for immunostaining as we have previously described^34^. Briefly, cells were stained for with alexa fluor 488 phalloidin (ThermoFisher Scientific) and subsequently for VE-Cadherin or heparan sulfate. The corresponding secondary antibody conjugated with alexa fluor 594 for 1 hour at room temperature. DAPI was applied for nuclear staining. For negative controls, samples were incubated with mouse IgG. Cells were then mounted with Aqua poly mount (Polysciences) and analyzed on a Leica TCS-SP2 AOBS confocal microscope (Leica Microsystems). All the images were taken with oil-immersed x40 objective, NA 1.25 and x63 objective, NA 1.4.

### 9. HPSE2 purification

HPSE2 was purified from conditioned medium of HEK 293T cells infected with HPSE2-overexpression lentivirus. Conditioned medium was collected, centrifuged at 3500 rpm for 10 min, and filtered through 0.22 μm filter. HPSE2 was collected with flow-through from CaptoQ ion exchange column equilibrated with 20 mmol/L Tris pH 8.7. HPSE2-containing fractions were collected, buffer was changed to 20 mmol/L MES pH 6.0 containing 100 mmol/L NaCl. HPSE2 was elute with 0.1 – 1 mol/L NaCl gradient. HPSE2 fractions purity was assessed by electrophoresis followed by gel staining with Colloidal Blue Staining Kit (ThermoFisher Scientific) and by western blotting. HPSE2 was collected, concentrated using centricons, and dialysed against PBS.

### 10. Animal experiments

Sepsis models in mice. All procedures were carried out at Phenos GmbH (Hannover, Germany) The animal protection committee of the local authorities (Lower Saxony state department for food safety and animal welfare [LAVES]) approved all experiments (approval: 33.19-42502-04-16/2255). The group size was determined using http://powerandsamplesize.com/ to achieve a detectable difference between different groups with an 80% probability at a 5% significance level. Eight-to 10-weeks-old male C57BL6 mice (20–25 g) obtained from Charles River (The Charles River Laboratories; Sulzfeld, Germany) were injected iv with purified HPSE2 (5μg/g body weight) and immediately thereafter low dose (5 ng/g body weight) of LPS (E.coli O111:B4, Sigma) administration was performed by iv injection. This LPS dose was chosen in preliminary experiments ranging from 5 to 1000 ng/ g body weight. 5 ng/g body weight LPS dose was well tolerated and resulted in a transient increase of pro-inflammatory cytokine level in blood within 2-4h after injection which declines at 8h and completely normalized at 20h post injection (data not shown). The blood sample were obtained at 2h post LPS injection, the plasma levels of pro-inflammatory cytokines IL-6 and TNFα were quantified by bead-based flow cytometry assay (CBA Kit; BD Biosciences, Heidelberg, Germany) according to the manufacturer’s instructions.

Polymicrobial sepsis in mice was induced by cecal ligation and puncture (CLP) as described previously^35^. Vehicle (PBS) or purified HPSE2 (100 μg in 50 μl) was administrated by intravenous injection 1h prior to CLP surgery. At 18 h after CLP or sham operation, mice were anesthetized with isofluorane for blood sampling. Subsequently animals were sacrificed and the kidneys were perfused with PBS solution via the left ventricle, removed, fixed in formalin for 24h, and processed for immunohistochemistry as described^35^.

### 11. Statistics

We used mean ± SEM throughout this study. All experiments were repeated independently at least three times. For comparing two different groups of data, Student’s T test test was applied. Multiple comparisons were analyzed by using the one-way analysis of variance with the Tukey as a post hoc test. “*” shows P value of less than 0.05; “**” shows P value of less than 0.01; “***” shows P value of less than 0.001. GraphPad Prism version 5.02 (GraphPad Prism Software Inc., San Diego, CA, USA) was used for data analysis.

## Results

### 1. HPSE1 aggravates endothelial LPS responses

We first treated cells with purified catalytically active HPSE1 and then stimulated the cells with LPS. Immunocytochemical staining using monoclonal 10E4 antibody confirmed removal of heparan sulfate structures by HPSE1 (Fig. 1A, B). Cells treated with HPSE1 demonstrated morphological changes similar to the changes induced by LPS, namely cytoskeletal rearrangement and disruption of cell-cell contacts (Fig. 1C, D). When HPSE1-treated endothelial cells were stimulated with LPS, expression of IL-6 was significantly higher than in untreated cells (Fig. 1E). IL-6 was selected as a readout of the LPS-induced TLR4 response because IL-6 expression in endothelial cells was shown to be TLR4-dependent^36^. In our experiments, the LPS-dependent expression of IL-6 was abrogated by the TLR4 inhibitor Cli-095 (data not shown). In a similar fashion, conditioned medium of the HPSE1-overexpressing endothelial cells that secrete the protein to the cell culture supernatant also promoted the LPS response in comparison to conditioned medium of the cells infected with control virus (Fig. 1F). This data showed that catalytic activity of HPSE1 promotes the LPS response in endothelial cells.

**Figure 1.**
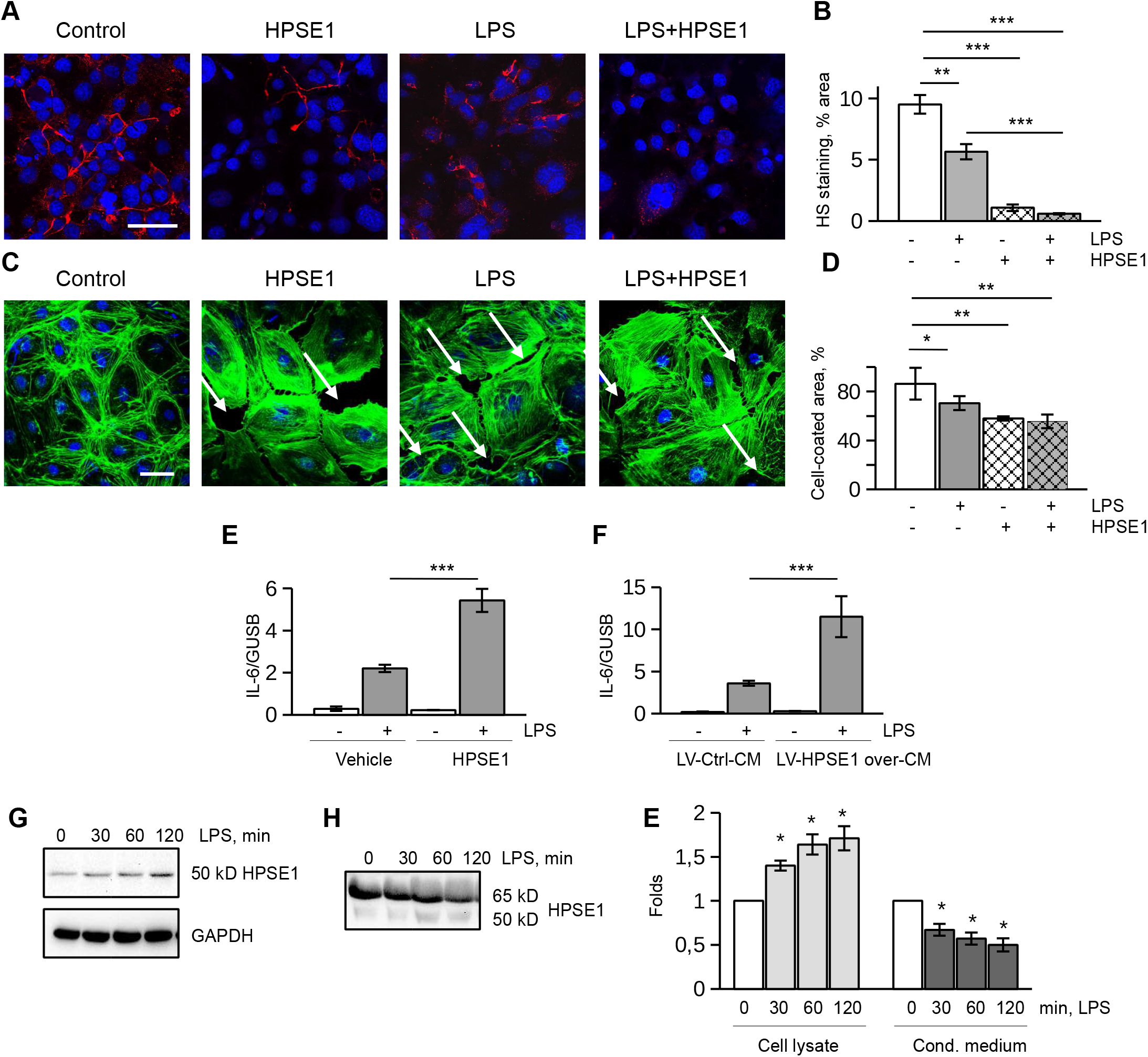
HPSE1 aggravates endothelial cell response to LPS. **A**. HMEC-1 cells were treated with active HPSE1 for 1 h prior to LPS stimulation. Then, cells were treated with 100 ng/ml LPS for 3 hrs. Cells were fixed and stained with 10A4 antibody for heparan sulfate (Alexa 594). Nuclei were stained with DAPI. Scale Bar 50μm **B**. Quantification of heparan sulfate stainings. Percentage of stained areas was calculated using ImageJ. **C.** Primary endothelial cells treated as in A were stained with Alexa 488-Phalloidin (green) and DAPI. White arrows show loss of cellular contacts. Scale Bar 50μm. **D.** Quantification of phalloidin images. Percentage of cell-coated areas was calculated using ImageJ. **E.** HMEC-1 cells were treated with catalytically active HPSE1 and then stimulated with LPS. IL-6 expression was assessed by RT-PCR. **F.** Cell were treated with conditioned medium from control (LV-Ctrl-CM) and HPSE1 overexpression (LV-HPSE1 over-CM) lentivirus-transduced HMEC-1 cells for 1h prior to LPS stimulation. IL-6 expression was assessed by RT-PCR. **G.** Expression of 50 kD active HPSE1 after LPS stimulation of endothelial cells assessed in cell lysates by western blotting. **H.** Expression of 65 kD and 50 kD isoforms of HPSE1 was assessed in conditioned medium of LPS-stimulated endothelial cells. **E.** Quantification of three independent western blotting experiments as shown in G (Cell lysate) and H (Cond. Medium). Data were quantified using QuantityOne software (BioRad).

Endothelial cells can serve as a source of HPSE1^26^. HPSE1 is expressed in the form of 65 kD inactive precursor that is processed to an active 50 kD isoform through cleavage by Cathepsin L (Ctsl)^6^. To assess whether or not endogenous HPSE1 is activated by LPS in endothelial cells, we analyzed the expression of both HPSE1 isoforms in the conditioned medium of cells and cell lysates using antibody that recognizes both forms. We found 65 kD inactive HPSE1 mainly in cell conditioned medium, whereas active 50 kD form was mostly associated with the cells. Furthermore, the content of the endogenous catalytically active 50 kD HPSE1 was time dependently increased after the LPS treatment (Fig. 1G-E) whereas the content of 65 kD isoform in the conditioned medium was decreasing. These data point that endogenous HPSE1 can be activated after LPS treatment to provide for a positive TLR4-signaling feedback loop.

### 2. HPSE2 overexpression protects from heparan sulfate-glycocalyx shedding

To visualize the role of both heparanases in the turnover of the heparan sulfate glycocalyx in the endothelial cells, we established overexpression of HPSE2 through lentiviral transduction (Suppl. Fig. S1A-C). Similar to the endogenous HPSE2, overexpressed protein was released from the cells and distributed between free soluble form in the conditioned medium and cellsurface bound form (Suppl. Fig. S1B, C). HPSE-2 overexpressing endothelial cells showed no signs of toxicity. Instead, proliferation rate was slightly increased and the cells were protected against the TGFβ-induced apoptosis (Suppl. Fig. S1D, E).

We next inspected HPSE2-overexpressing endothelial cells that were incubated in the Polydimethylsiloxan (PDMS) – microfluidic chips under medium flow conditions. The chips consisted of 4 parallel 10 mm-long and 2 mm-wide channels (Fig. 2A). The cells were incubated under permanent medium flow conditions for 3 days. Endothelial cells developed a more 3D heparan sulfate-enriched glycocalyx layer when were subjected to the medium flow (Fig. 2B). Orthogonal views of the 3D z-scans confirmed extracellular localization of the glycocalyx layer above the layer of cell nuclei (Fig. 2B). We then added catalytically active HPSE1 to the medium used for chips perfusion. Under flow conditions this treatment also led to the shedding of heparan sulfate layer from the cell surface (Fig. 2C, D). Remarkably, cells overexpressing HPSE2 were protected against glycocalyx shedding by the exogenous HPSE1 (Fig. 2C, D). Similar results were obtained when shear stress was increased up to 38.75 dyn/cm^2^ (Supp. Fig. S2A).

**Figure 2.**
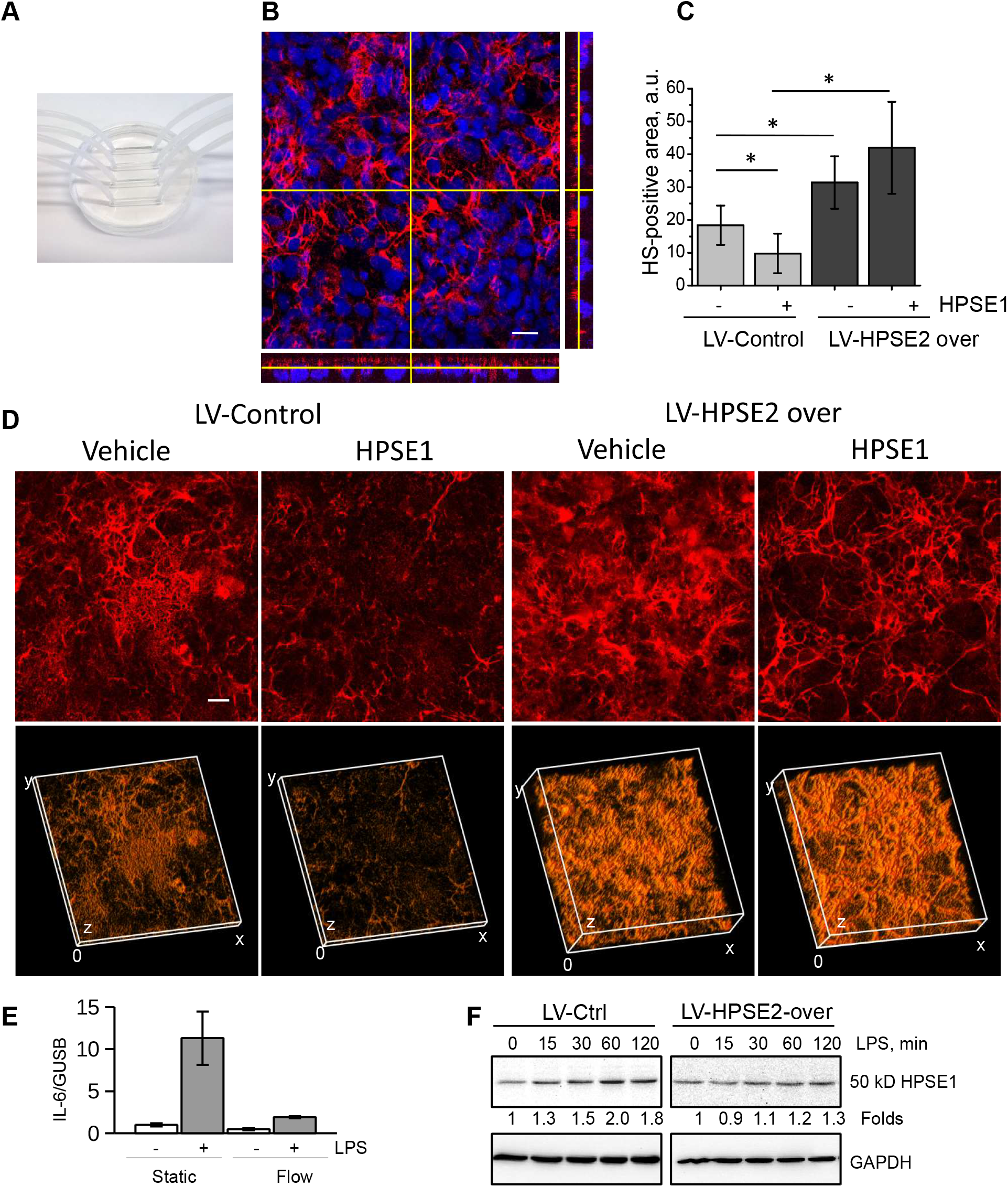
HPSE2 overexpression protects endothelial heparan sulfate glycocalyx in a microfluidic chip model. **A.** 4-channels PDMS-microfluidic chip. **B.** Development of 3D heparan sulfate glycocalyx layer after endothelial cells cultivation under flow conditions (3 days, 0.1 dyn/cm^2^) was assessed by 3D confocal microscopy using 10E4 antibody to heparan sulfate followed by image reconstruction using ImageJ software. DraQ5 was used as a nuclear stain. Yellow lines show position of orthogonal slices. Bottom and right panels show corresponding orthogonal slices of the sum images. **C.** Control and HPSE2 overexpression-lentivirus-infected endothelial cells were cultivated in the microfluidic chip for 3 days, and then perfused with recombinant HPSE1 for 4 hrs. Cells were fixed by perfusion with PFA as described in the methods section. Heparan sulfate content was quantified using ImageJ software after 3D confocal microscopy. Experiment was independently repeated in 4 chips. **D.** 3D reconstruction of heparan sulfate glycocalyx layer images of lentivirus-infected endothelial cells after incubation in the microfluidic chip and recombinant HPSE1 treatment. The 3D reconstruction was performed using ImageJ software. Scale bar 20μm. **E.** Endothelial cells incubated in microfluidic chips under static and flow conditions were stimulated with 100ng/ml LPS for 3 hrs. IL-6 expression was assessed by TaqMan RT-PCR. **F.** 50 kD HPSE1 expression in lysates of LPS-stimulated endothelial cells.

Confirming inhibitory role of heparan sulfate on the TLR4 response, the cells cultured under flow conditions demonstrated strongly diminished response to LPS (Fig. 2E). The expression of HPSE1 in endothelial cells cultured under flow was also diminished (Suppl. Fig. S2B).

To assess whether or not HPSE2 prevents also the activity of endogenous HPSE1, lentivirus-infected cells were stimulated with LPS and activation of HPSE1 was followed by the accumulation of the active 50 kD HPSE1 isoform (Fig. 2F, Suppl. Fig. S1F). The overexpression of HPSE2 not only prevented the activity of exogenous HPSE1, but also LPS-dependent increase of endogenous HPSE1 activity. This data confirmed antagonistic relationship between the heparanases in endothelial cells. Accordingly, both expression and activity of Ctsl in the HPSE2 overexpressing cells were slightly decreased (Supp. Fig. S2B, C).

### 3. HPSE2 protects endothelial cells

We next studied whether or not HPSE2 can also exert protective functions under pathological situations. Control and HPSE2-overexpressing endothelial cells cultured under static conditions were treated with different concentrations of LPS. Significant downregulation of the LPS-induced IL-6 expression was observed in the HPSE2-overexpressing endothelial cells at different LPS concentrations (Fig. 3A). To precisely characterize the LPS response in HPSE2 overexpressing cells, Human Common Cytokine RT^2^ Profiler Array was performed. Expressions of several pro-inflammatory cytokines, in particular, IL-6, IFNB1, IFNA1, CSF3 (G-CSF), and TNFSF10 were significantly downregulated in HPSE2-overexpressing cells (Suppl. Fig. S3A). The data of the array were verified in independent RT-PCR experiment (Suppl. Fig. S3B-E). Further cytokines including CSF2, IL-1A, IL-1B, TGFB3, TNFRSF11 were downregulated to a lesser degree. Together, these data demonstrated general anti-inflammatory effects of the HPSE2 overexpression in the endothelial cells. The overexpression of HPSE2 have not induced any inflammatory effect on endothelial cells.

**Figure 3.**
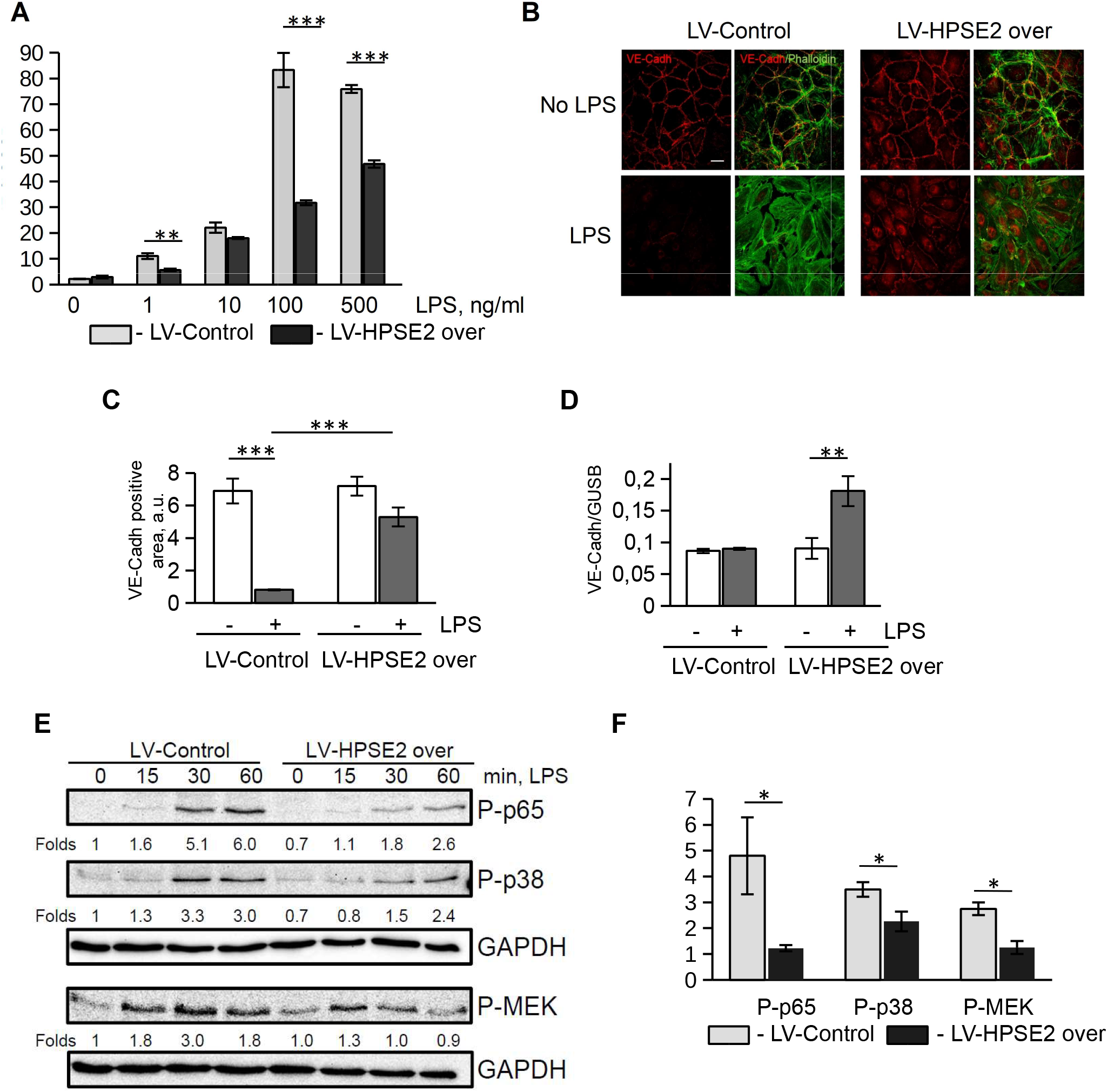
HPSE2 protects endothelial cells from LPS-induced damage. **A.** Endothelial cells infected with Control and HPSE2-overexpression lentivirus were treated with different concentrations of LPS for 3 hrs as indicated. IL-6 expression was assessed by TaqMan RT-PCR. **B.** Primary endothelial cells treated with LPS for 24 hrs cells were fixed and immunostained for VE-Cadherin (Alexa 594) and Phalloidin (Alexa 488). **C.** The percentage of VE-Cadherin positive areas was calculated using ImageJ. **D.** Expression of VE-Cadherin mRNA in LPS treated cells was quantified by TaqMan RT-PCR. **E.** Endothelial cells were stimulated with 1 μg/ml LPS for indicated times. Protein phsophorylation was assessed by western blotting. **F.** Quantification of phosphorylation of p65, p38, p-MEK at 30 min of LPS stimulation quantified from three independent western blotting experiments.

We then assessed morphological changes in the endothelial cells treated with LPS. Similar to data reported by Zheng et al.^37^, we observed that control cells treated with 100 ng/ml LPS lost cell-cell contacts and demonstrated decreased VE-Cadherin protein expression (Fig. 3B, C), while HPSE2-overexpressing cells were protected and preserved cell-cell contacts much better than control cells. These cells also demonstrated increased VE-Cadherin expression after the LPS treatment (Fig. 3D). LPS-induced protein phosphorylation and activation of NFκB were less activated in HPSE-2 overexpressing endothelial cells as determined by western blotting (Fig. 3E, F) and luciferase assay (Suppl. Fig. S2E). Together, these data demonstrated a strong protection of endothelial cells exposed to LPS by HPSE2.

### 4. HPSE2 prevents HPSE1 activity

To test this idea, we made use of OGT 2115, an inhibitor of HPSE1 catalytic activity. The compound decreased NFκB activation as shown in promoter activity luciferase assay (Suppl. Fig. S2E). LPS-dependent expression and secretion of IL-6 was accordingly downregulated in the presence of HPSE1 inhibitor (Fig. 4A, B). However, there was no additive effect after HPSE1 inhibition in HPSE2 overexpressing cells (Fig. 4A, B; Suppl. Fig. S2E). This finding suggested that both proteins are parts of the same molecular pathway and anti-inflammatory role of HPSE2 is most likely mediated by its antagonistic action on HPSE1. A specific protective role of HPSE2 was further confirmed using the 1c7 antibody to HPSE2^23^ that can block HPSE2 interaction with heparan sulfate^38^. Application of the 1c7 antibody and thus displacement of the protein from cell surface heparan sulfate glycocalyx structures resulted in abrogation of HPSE2 protective action and strong activation of LPS response (Fig. 4C). We suggest that HPSE1regulates TLR4 response in endothelial cells. The effects are counteracted by HPSE2 bound to cell surface heparan sulfate.

**Figure 4.**
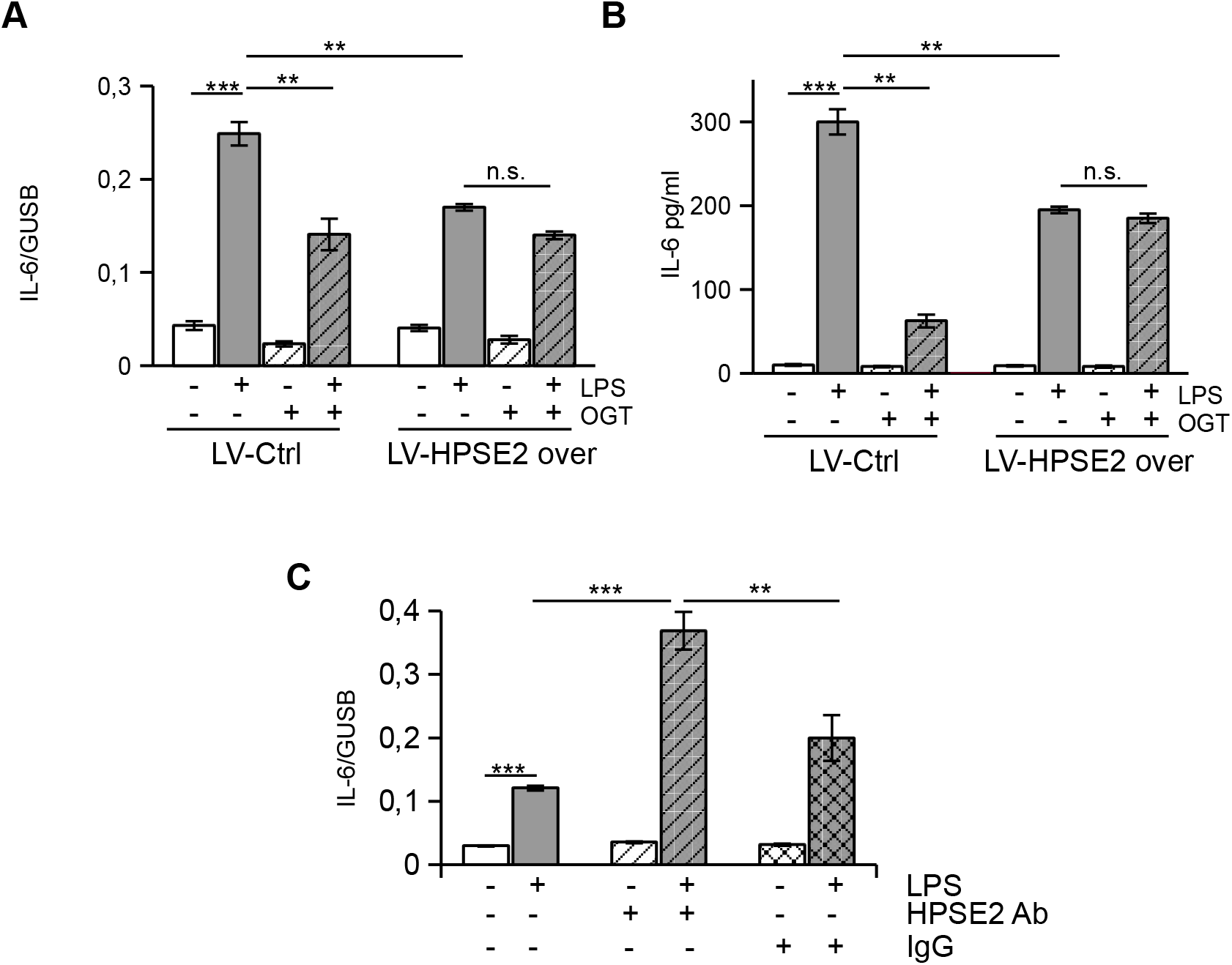
HPSE2 prrevents HPSE1 activity. **A.** Endothelial cells were pre-treated with inhibitor of HPSE1 catalytic activity, OGT 2115, and then stimulated with 100 ng/ml LPS for 3 hrs (A) and overnight (B). IL-6 expression was analyzed by RT-PCR (A) and ELISA (B). **C.** Endothelial cells were pre-incubated with mouse IgG or anti-HPSE2 antibody 1c7 for 30 min prior to LPS stimulation and then stimulated with 100 ng/ml LPS for 3 hrs. IL-6 expression was assessed by RT-PCR.

### 5. HPSE2 inhibits LPS binding to TLR4 receptor complex

We next studied the mechanisms of the regulation of LPS response in endothelial cells by HPSE1. Molecular mechanisms of TLR4 signaling are extensively investigated. The LPS molecule is first bound to TLR4 co-receptor, CD14^39^. Thereafter, LPS is transferred to the complex of TLR4 with an accessory molecule, Lymphocyte antigen 96, (also known as MD2) to induce the activation of intracellular adapters and subsequent protein phosphorylation cascades^40^. Several mechanisms could account for the role of HPSE1. First, TLR4 can be further activated by heparan sulfate fragments originating from the HPSE1 catalytic activity^16^. Second, LPS ligand binding to the CD14 receptor could be affected by heparan sulfate. Finally, the interaction of TLR4 with co-receptors on the cell membrane can be regulated by heparan sulfate. We first tested whether accumulation of heparan sulfate fragments during the cells stimulation with LPS can lead to further activation of TLR4. Exogenous heparan sulfate caused a weak increase in IL-6 expression, even at high concentrations (Fig. 5A). This response was also decreased in HPSE2-overexpressing cells (Fig. 5A). To investigate whether or not endogenously produced soluble heparan sulfate fragments can promote TLR4 activation, conditioned medium of cells treated with LPS for 3 h, was used for stimulation of naïve cells. We found that supernatant of LPS-treated cells caused strong increase of IL-6 expression (Fig. 5B). Proteinase K is a serine protease used to digest proteins in biological samples. Removal of protein components from the supernatant by incubation with beads-immobilized proteinase K prevented expression of IL-6. Removal of remaining LPS using Endotoxin-Removal beads caused complete abrogation of IL-6 expression (Fig. 5B). Exogenous heparan sulfate at concentration of 100 ng/ml strongly potentiated endothelial cell response to LPS (Fig. 5C). This response was documented by the 3-folds increase of the LPS-induced IL-6 expression in the presence of exogenous heparan sulfate. These findings suggested that concentration of endogenous heparan sulfate fragments produced during 3 hrs of LPS stimulation was not sufficient to potentiate activation of TLR4. However, in the case of significant accumulation, soluble heparan sulfate fragments can potentiate inflammatory response of endothelial cells to LPS.

**Figure 5.**
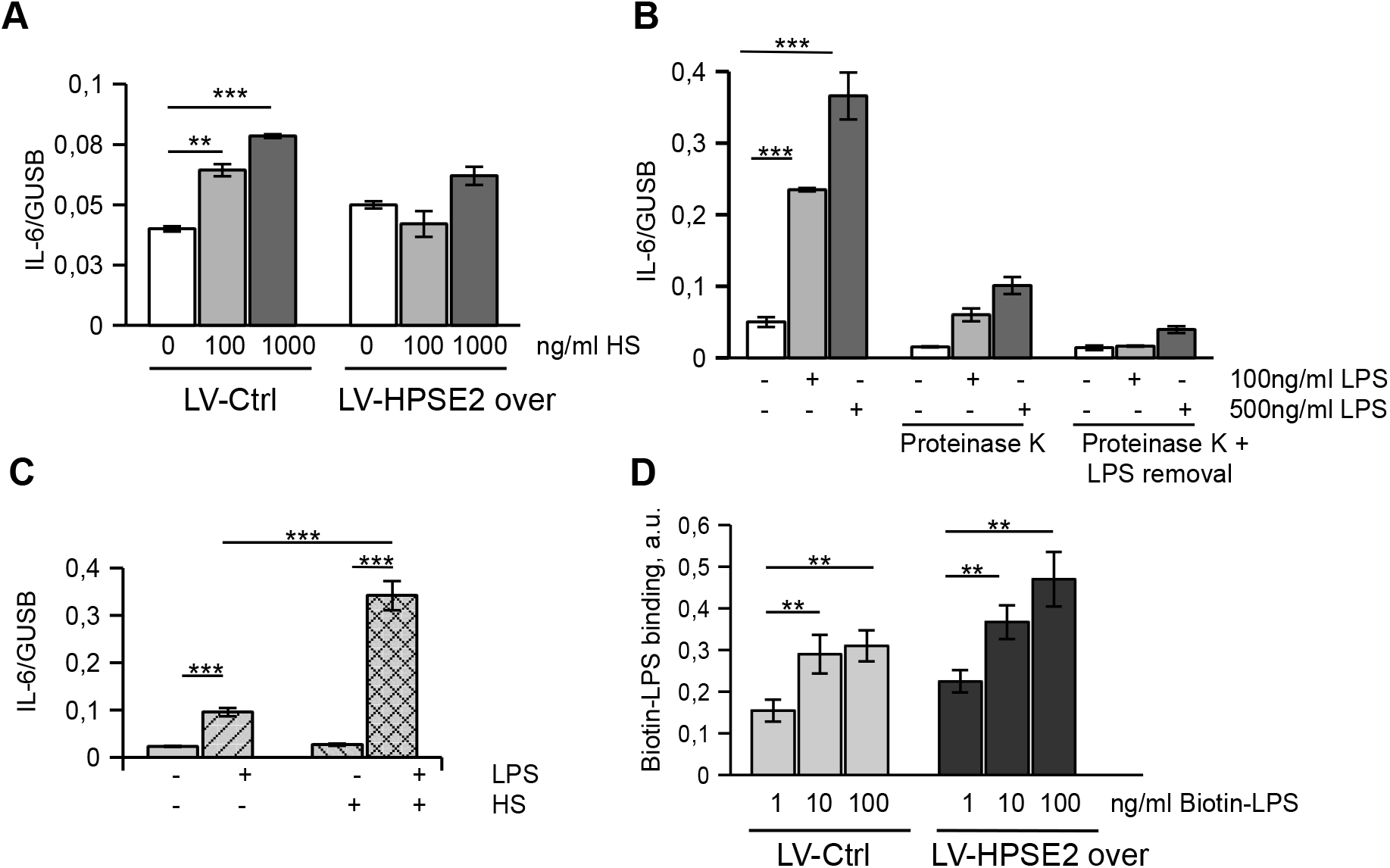
Heparan sulfate fragments cause weak activation of endothelial cells. **A.** Endothelial cells were treated with the indicated concentration of heparan sulfate (HS) for 3 hrs. IL-6 expression was assessed by RT-PCR. **B.** Conditioned medium of endothelial cells treated for 3 hrs with 100 ng/ml and 500 ng/ml LPS was used for stimulating naïve cells. Conditioned medium was treated with Proteinase K immobilized on agarose beads and remaining LPS was removed by incubation with LPS removal beads, as indicated. IL-6 expression was assessed by RT-PCR. **C.** Endothelial cells were stimulated with 100 ng/ml LPS in the absence and in the presence of 100 ng/ml heparan sulfate (HS). **D.** Cell-based ELISA for Biotin-LPS binding. Lentivirus-transduced endothelial cells were incubated on ice for 1h in the presence of biotin-LPS as indicated. After washing cells were incubated with SV-HRP. Biotin-LPS binding was quantified using TMB substrate kit.

We then investigated whether HPSE2 can affect LPS binding using biotin-LPS. Cells were incubated on ice to prevent internalization of receptors. Cell binding of biotin-LPS was not decreased by overexpression of HPSE2 (Fig. 5D), suggesting that LPS binding to CD14 was not affected. The specificity of biotin-LPS binding to CD14 was confirmed by application of CD14 blocking antibody. Over 70% of the Biotin-LPS binding was blocked by CD14 antibody but not isotypic IgG.

TLR4 intracellular signaling is mediated by the recruitment of two adapter proteins, MyD88 and TIR-domain-containing adapter-inducing interferon-β (TRIF)^41^. We observed diminished LPS-induced expression of IL-6 suggesting impaired MyD88-pathway activation in HPSE2-overexpressing cells. We further investigated the expression of IFNB1 and chemokine ligand-5 (CCL5), also known as RANTES that is mediated by TRIF adapter pathway activation (Fig. 6A). Our data show that both, MyD88 and TRIF-mediated signaling pathways of TLR4 are inhibited by HPSE2. Since LPS binding was not impaired by the overexpression of HPSE2, we reasoned that HPSE2 most likely interfered with the LPS transfer from CD14 to TLR4/MD2 complex. In order to verify this hypothesis, a pull-down assay using biotin-LPS and streptavidin magnetic beads was performed. Cells were incubated for 1 h on ice in the presence of 50 ng/ml biotin-LPS to allow its binding to the cell surface. Then, the cells were transferred to 37°C for the indicated time to enable further signaling events to take place. The results supported our hypothesis (Fig. 6B, C) as less TLR4 was detected in the biotin-LPS complexes from HPSE2-overexpressing endothelial cells. Regulation of CD14/TLR4 association by HPSE1 and protective role of HPSE2 was confirmed by the Duolink proximity ligation assay – a method to detect protein-protein interaction (Fig. 6D, E). To investigate the endocytosis of TLR4, cell surface staining on living cells was performed. To stop endocytosis and prevent staining of intracellular TLR4, cells were placed on ice after LPS stimulation and stained with anti-TLR4 antibody (Fig. 6F, G). Decreased staining suggested active endocytosis of the receptor complex in control cells. However, in HPSE2-overexpressing cells TLR4 remained on the cell surface. These findings imply that membrane-bound heparan sulfate prevents LPS transfer from CD14 to TLR4/MD2 complex and its removal by HPSE1 promotes the LPS response of endothelial cells.

**Figure 6.**
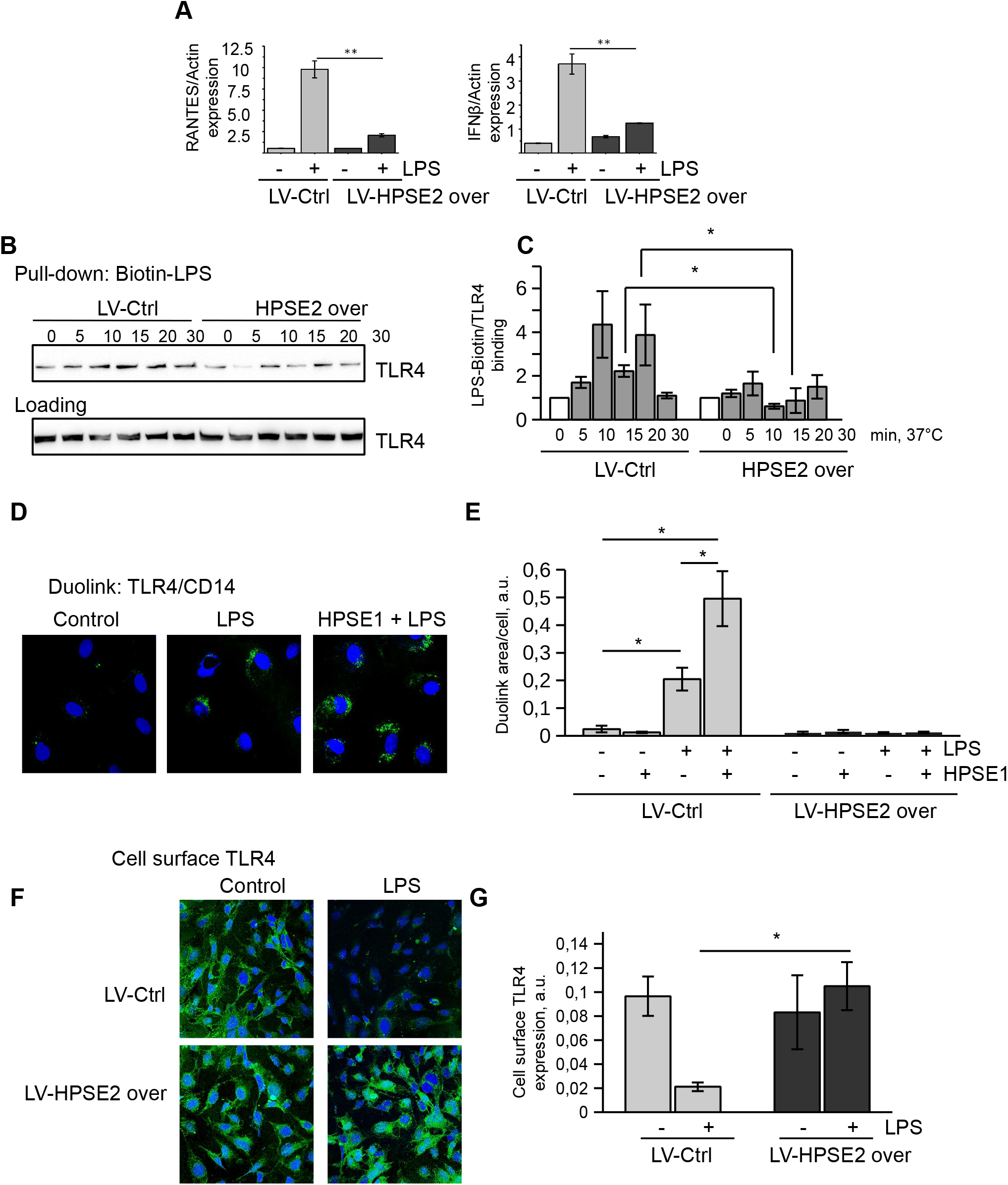
HPSE2 overexpression prevents TLR4 activation by LPS. **A.** Endothelial cells infected with Control and HPSE2 overexpression lentivirus were treated with 100 ng/ml LPS for 3 hrs. Expression was analyzed by TaqMan RT-PCR. **B.** TLR4 pull-down using biotin-LPS was performed as described in the Materials and Methods section. TLR4 was detected by immunoblotting. **C.** Quantification of independent TLR4 pull-down assay experiments. **D.** Duolink proximity ligation assay revealing direct interaction of TLR4 and CD14 was performed on endothelial cells stimulated with 100 ng/ml LPS for 20 min. Treatment with catalytically active HPSE1 was performed for 1 h prior to LPS stimulation. **E.** Duolink signal stained as in F was quantified using ImageJ as described in Materials and Methods. **F.** To detect cell surface TLR4 lentivirus-infected endothelial cells were stimulated with 500 ng/ml LPS for 1h, then placed on ice and incubated with TLR4 antibody. Then, the cells were washed, fixed, and stained with secondary antibody (green) and DAPI. **G.** Cell surface TLR4 expression of lentivirus-infected and LPS-stimulated endothelial cells was quantified using cell-based ELISA as described in Materials and Methods.

### 6. HPSE2 is protective in vivo

To investigate how expression of endogenous HPSE2 is regulated during sepsis, we used the mouse CLP polymicrobial sepsis model. We observed decreased HPSE2 expression in serum of CLP mice (Fig. 7A, B). In the kidney we observed HPSE2 expression in medullary capillaries. This expression of HPSE2 was strongly downregulated in sepsis (Fig. 7C, D). Negative control staining is shown in supplementary Fig. S3F.

**Figure 7.**
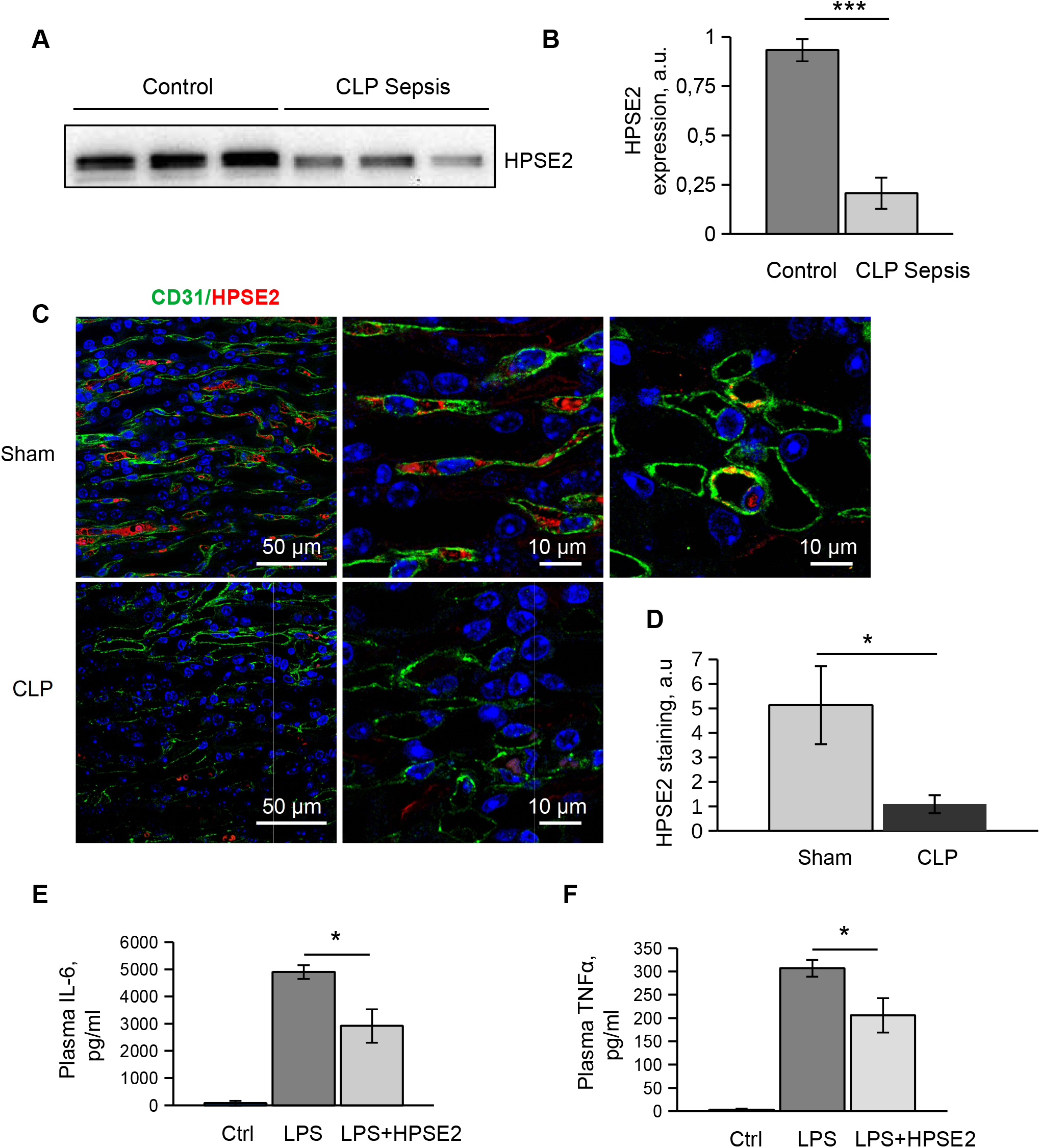
HPSE2 protects against inflammation in vivo. **A.** Serum level of HPSE2 expression in mouse CLP sepsis model was assessed by western blotting. **B.** Qantification of westernblotting data. **C.** Expression of HPSE2 in mouse kidney medullary capillaries in CLP sepsis model. Tissue sections were stained with CD31(Alexa 488) and HPSE2 (Alexa 594). Nuclei were stained with DAPI. **D.** Quantification of HPSE2 immunohistochemistry stainings in mouse kidney. **E, F.** Mice were iv injected with 5μg/g body weight LPS alone or LPS alone with 5μg/g body weight purified HPSE2. After 2hrs plasma cytokine level was measured using cytometric bead array. (E) IL-6 expression; (F) TNFα expression. Mean ±s.e.m. is shown.

We next purified HPSE2 from conditioned medium of HEK293T cells transduced with LV-HPSE2 overexpression virus. Application of the exogenous HPSE2 diminished LPS response of the endothelial cells in vitro in a similar fashion to HPSE2 overexpression (Supplementary Fig. S3G). When co-injected intravenously along with LPS in mice, HPSE2 had similar protective effects and caused statistically significant decrease in the plasma content of TNFα and IL-6 2 h after injection (Fig. 7E, F). This data suggest that HPSE2 has also an endothelial protective function in microvasculature in vivo and can have some therapeutic potential.

## Discussion

We demonstrated an important protective role for HPSE2 both in vivo and in vitro. Using different approaches we showed that HPSE1 activity aggravates endothelial response to LPS, whereas HPSE2 prevents activation of HPSE1, heparan sulfate shedding, and inflammatory responses mediated by TLR4. Our findings support an essential role for an intact glycocalyx in the endothelium. Glycocalyceal shedding occurs in many diseases and pathological insults.

However, the mechanisms how glycocalyceal turnover is regulated and what the consequences are when the process is perturbed is still imperfectly understood. A high expression of HPSE1, the sole enzyme capable of cleaving heparan sulfate, is damaging for the endothelium^42^. Although the role of HPSE2 as an antagonist of HPSE1 is well appreciated in cancer, how the interaction of these proteins can affect endothelial cells is less clear.

The role of heparan sulfate in the TLR4-mediated LPS response is not well understood. Soluble heparan sulfate fragments generated by HPSE1 can serve as ligand for TLR4^16^. Treatment of macrophages with HPSE1 facilitated an inflammatory response. On the contrary, in microglia heparan sulfate promoted TLR4/CD14 complex formation and HPSE1 transgenic cells showed less inflammatory response to LPS^20^. Our data suggest that heparan sulfate prevents the activation of TLR4 signaling and inflammatory response of endothelium. HPSE1 is an important component of the TLR4 signalosome as heparan sulfate cleavage facilitates cellular response and promotes inflammatory reaction of the endothelial cells.

Both heparanases are present in human plasma. HPSE1 upregulation is associated with many disease states. Our data showed decreased plasma HPSE2 expression in mouse sepsis models indicating that cross-talk of the heparanases is important in vivo. Our data suggest that endothelial cell express and release HPSE1 and HPSE2 for local regulation of heparan sulfate turnover. Our mechanistic studies showed that HPSE2 bound to heparan sulfate on the endothelial cell surface prevents activation of HPSE1 and heparan sulfate shedding. We found that HPSE2 displacement from the cell surface by the 1c7 antibody^38^ led to the abrogation of HPSE2 protective function. Taking into account poor development of the heparan sulfate glycocalyx under normal static cell culture conditions we applied microfluidic chip model to demonstrate the protective role of HPSE2^43,44^. In the microfluidic experiments, HPSE2 overexpressing cells were strongly protected from heparan sulfate loss by HPSE1. This observation further confirmed that HPSE2 expressed by endothelial cells partially remained bound to the heparan sulfate glycocalyx and counteracted its cleavage by HPSE1. Inhibiting role of the glycocalyx in TLR4 receptor signaling was confirmed by diminished LPS response of the cells incubated under flow conditions.

In macrophages HPSE1 itself was shown to induce TLR dependent signaling^21^. In endothelial cells HPSE1 induced rather weak inflammatory response by itself and only by the prolonged incubation (data not shown). Thus, in the context of TLR4 activation on the endothelium, remodeling of the glycocalyx on the cell surface seems more important. Interestingly, exogenous heparan sulfate strongly potentiated cell response to LPS. The finding can probably be explained by competitive binding of HPSE2 to soluble heparan sulfate resulting in its’ removal from the cell membrane-associated glycocalyx.

We used biotin-LPS to demonstrate that HPSE2 does not interfere with LPS binding to the CD14 but rather prevents LPS transfer to TLR4 and activation of the cellular signaling. As a result, the inflammatory response was reduced. Early reports suggested that endothelial cells express only the soluble form of CD14^45,46^. On the contrary, more recent reports showed the expression and the role for membrane CD14 in endothelial response to LPS^47,48^. Lloyd-Jones et al. showed that membrane-bound CD14 is indispensable for the activation of TRIF-mediated pathway in endothelial cells^48^. We showed that both, MyD88- and TRIF-dependent pathways are affected by HPSE2. Therefore, we assume that membrane bound form of CD14 is regulated by heparan sulfate. However, involvement of soluble CD14 can not be excluded. In vivo co-injection of HPSE2 along with LPS in mice led to decreased cytokines expression in plasma. In polymicrobial CLP mouse sepsis model the expression of HPSE2 in renal medullary capillaries was decreased. Thus, loss of HPSE2 can promote microcirculatory disorder in sepsis leading to organ failure.

Our results suggest that HPSE2, expressed locally by the endothelial cells or delivered with blood, fulfills protective role in microvasculature via protection from heparan sulfate shedding and anti-inflammatory regulation of TLR4 signaling. HPSE2 is a novel molecule which exerts a direct protective effect on the endothelial glycocalyx thereby maintaining microvascular function and stability as well as protecting the endothelium from damage. Our results in vivo suggest that HPSE2 supplementation may be beneficial for the protection of the microvasculature. This novel mechanism supports therapeutic strategies to stabilize the endothelial glycocalyx.

## Acknowledgments

a. We are grateful to Prof. Israel Vlodavsky (Technion, Haifa, Israel) for giving us 1c7 antibody to HPSE2.
b. Sources of funding: Grants from German Federal Ministry of Education and Research (BMBF) Nr. 031A577A and 031A577B funded this research. This work was also supported by a grant for the German Research Council to H.H. Ha 1388/17-1.
c. Disclosures: none.

## Highlights

➢ Catalytic activity of Heparanase 1 aggravates TLR4 signaling and endothelial response to LPS
➢ Heparanase 2 diminishes endothelial response to LPS
➢ Heparanase 2 does not affect LPS binding to CD14 but inhibits LPS presentation to TLR4
➢ Expression of HPSE2 in vivo is decreased in sepsis.

## References

1. Weinbaum S, Tarbell JM, Damiano ER. The Structure and Function of the Endothelial Glycocalyx Layer. Annual Review of Biomedical Engineering. 2007;9(1):121–167.

2. Schmidt EP, Yang Y, Janssen WJ, Gandjeva A, Perez MJ, Barthel L, Zemans RL, Bowman JC, Koyanagi DE, Yunt ZX, Smith LP, Cheng SS, Overdier KH, Thompson KR, Geraci MW, et al. The pulmonary endothelial glycocalyx regulates neutrophil adhesion and lung injury during experimental sepsis. Nature Medicine. 2012;18(8):1217–1223.

3. Florian JA. Heparan Sulfate Proteoglycan Is a Mechanosensor on Endothelial Cells. Circulation Research. 2003;93(10):136e–142.

4. Ori A, Wilkinson MC, Fernig DG. A systems biology approach for the investigation of the heparin/heparan sulfate interactome. Journal of Biological Chemistry. 2011;286(22):19892–19904.

5. Ushiyama A, Kataoka H, Iijima T. Glycocalyx and its involvement in clinical pathophysiologies. Journal of Intensive Care. 2016;4(1): 1–11.

6. Rabelink TJ, de Zeeuw D. The glycocalyx—linking albuminuria with renal and cardiovascular disease. Nature Reviews Nephrology. 2015; 11(11):667–676.

7. Kim YH, Nijst P, Kiefer K, Tang WHW. Endothelial Glycocalyx as Biomarker for Cardiovascular Diseases: Mechanistic and Clinical Implications. Current Heart Failure Reports. 2017;14(2):117–126.

8. Song JW, Zullo J, Lipphardt M, Dragovich M, Zhang FX, Fu B, Goligorsky MS. Endothelial glycocalyx-the battleground for complications of sepsis and kidney injury. Nephrology Dialysis Transplantation. 2018;33(2):203–211.

9. Chelazzi C, Villa G, Mancinelli P, De Gaudio AR, Adembri C. Glycocalyx and sepsis-induced alterations in vascular permeability. Critical Care. 2015;19(1):1–7.

10. Martin L, Koczera P, Zechendorf E, Schuerholz T. The Endothelial Glycocalyx: New Diagnostic and Therapeutic Approaches in Sepsis. BioMed Research International. 2016;2016.

11. Lin X, Wei G, Shi Z, Dryer L, Esko JD, Wells DE, Matzuk MM. Disruption of gastrulation and heparan sulfate biosynthesis in EXT1-deficient mice. Developmental Biology. 2000;224(2):299–311.

12. Stickens D. Mice deficient in Ext2 lack heparan sulfate and develop exostoses. Development. 2005;132(22):5055–5068.

13. Melo CM, Origassa CST, Theodoro TR, Matos LL, Miranda TA, Accardo CM, Bouças RI, Suarez ER, Pares MMNS, Waisberg DR, Toloi GC, Nader HB, Waisberg J, Pinhal MAS. Análises das isoformas de heparanase e da catepsina B em plasma de pacientes com carcinomas gastrointestinais: Estudo transversal analítico. Sao Paulo Medical Journal. 2015;133(1):28–35.

14. Shafat I, Ilan N, Zoabi S, Vlodavsky I, Nakhoul F. Heparanase levels are elevated in the urine and plasma of type 2 diabetes patients and associate with blood glucose levels. PLoS ONE. 2011;6(2).

15. Vijay K. Toll-like receptors in immunity and inflammatory diseases: Past, present, and future. International Immunopharmacology. 2018;59(February):391–412.

16. Goodall KJ, Poon IKH, Phipps S, Hulett MD. Soluble heparan sulfate fragments generated by heparanase trigger the release of pro-inflammatory cytokines through TLR-4. PLoS ONE. 2014;9(10).

17. Akbarshahi H, Axelsson JBF, Said K, Malmström A, Fischer H, Andersson R. TLR4 dependent heparan sulphate-induced pancreatic inflammatory response is IRF3-mediated. Journal of Translational Medicine. 2011;9(1): 1–8.

18. Lerner I, Hermano E, Zcharia E, Rodkin D, Bulvik R, Doviner V, Rubinstein AM, Ishai-Michaeli R, Atzmon R, Sherman Y, Meirovitz A, Peretz T, Vlodavsky I, Elkin M. Heparanase powers a chronic inflammatory circuit that promotes colitis-associated tumorigenesis in mice. Journal of Clinical Investigation. 2011;121(5): 1709–1721.

19. Brunn GJ, Bungum MK, Johnson GB, Platt JL. Conditional signaling by Toll-like receptor 4. The FASEB Journal. 2005;19(7):872–874.

20. O’Callaghan P, Li JP, Lannfelt L, Lindahl U, Zhang X. Microglial heparan sulfate proteoglycans facilitate the cluster-of-differentiation 14 (CD14)(WARNING)Toll-like receptor 4 (TLR4)-dependent inflammatory response. Journal of Biological Chemistry. 2015;290(24):14904–14914.

21. Blich M, Golan A, Arvatz G, Sebbag A, Shafat I, Sabo E, Cohen-Kaplan V, Petcherski S, Avniel-Polak S, Eitan A, Hammerman H, Aronson D, Axelman E, Ilan N, Nussbaum G, et al. Macrophage activation by heparanase is mediated by TLR-2 and TLR-4 and associates with plaque progression. Arteriosclerosis, Thrombosis, and Vascular Biology. 2013;33(2).

22. McKenzie E, Tyson K, Stamps A, Smith P, Turner P, Barry R, Hircock M, Patel S, Barry E, Stubberfield C, Terrett J, Page M. Cloning and expression profiling of Hpa2, a novel mammalian heparanase family member. Biochemical and Biophysical Research Communications. 2000;276(3):1170–1177.

23. Levy-Adam F, Feld S, Cohen-Kaplan V, Shteingauz A, Gross M, Arvatz G, Naroditsky I, Ilan N, Doweck I, Vlodavsky I. Heparanase 2 interacts with heparan sulfate with high affinity and inhibits heparanase activity. Journal of Biological Chemistry. 2010;285(36):28010–28019.

24. Vlodavsky I, Gross-cohen M, Weissmann M, Ilan N, Ralph D. Opposing functions of heparanase-1 and heparanase-2 in cancer progression. Trends Biochem Sci. 2018;43(1): 18–31.

25. Chen G, Wang D, Vikramadithyan R, Yagyu H, Saxena U, Pillarisetti S, Goldberg IJ. Inflammatory Cytokines and Fatty Acids Regulate Endothelial Cell Heparanase Expression. Biochemistry. 2004;43(17):4971–4977.

26. Godder K, Vlodavsky I, Eldor A, Weksler BB, Haimovitz-Freidman A, Fuks Z. Heparanase activity in cultured endothelial cells. Journal of Cellular Physiology. 1991;148(2):274–280.

27. Edovitsky E, Lerner I, Zcharia E, Peretz T, Vlodavsky I, Elkin M. Role of endothelial heparanase in delayed-type hypersensitivity. Blood. 2006;107(9):3609–3616.

28. Hodjat M, Haller H, Dumler I, Kiyan Y. Urokinase receptor mediates doxorubicin-induced vascular smooth muscle cell senescence via proteasomal degradation of TRF2. Journal of Vascular Research. 2013;50(2).

29. Zufferey R, Nagy D, Mandel RJ, Naldini L, Trono D. Multiply attenuated lentiviral vector achieves efficient gene delivery in vivo. Nature Biotechnology. 1997;15(9):871–875.

30. Naldini L, Blomer U, Gallay P, Ory D, Mulligan R, Gage FH, Verma IM, Trono D. In Vivo Gene Delivery and Stable Transduction of Nondividing Cells by a Lentiviral Vector. Science. 1996;272(5259):263–267.

31. Farsari M, Chichkov BN. Materials processing: Two-photon fabrication. Nature Photonics. 2009;3(8):450–452.

32. Duffy DC, McDonald JC, Schueller OJA, Whitesides GM. Rapid prototyping of microfluidic systems in poly(dimethylsiloxane). Analytical Chemistry. 1998;70(23):4974–4984.

33. Schappe MS, Desai BN. Measurement of TLR4 and CD14 Receptor Endocytosis Using Flow Cytometry. Bio-protocol. 2018;8(14):e2926.

34. Kiyan J, Kiyan R, Haller H, Dumler I. Urokinase-induced signaling in human vascular smooth muscle cells is mediated by PDGFR-β. EMBO Journal. 2005;24(10).

35. Menne J, Shushakova N, Bartels J, Kiyan Y, Laudeley R, Haller H, Park J-K, Meier M. Dual inhibition of classical protein kinase C-α and protein kinase C-ß isoforms protects against experimental murine diabetic nephropathy. Diabetes. 2013;62(4).

36. Anjum SA, Lawrence H, Holland JP, Kirby JA, Deehan DJ, Tyson-Capper AJ. Effect of cobalt-mediated Toll-like receptor 4 activation on inflammatory responses in endothelial cells. Oncotarget. 2016;7(47):1–8.

37. Zheng X, Zhang W, Hu X. Different concentrations of lipopolysaccharide regulate barrier function through the PI3K/Akt signalling pathway in human pulmonary microvascular endothelial cells. Scientific Reports. 2018;8(1):1–11.

38. Gross-Cohen M, Feld S, Doweck I, Neufeld G, Hasson P, Arvatz G, Barash U, Naroditsky I, Ilan N, Vlodavsky I. Heparanase 2 attenuates head and neck tumor vascularity and growth. Cancer Research. 2016;76(9):2791–2801.

39. Ryu JK, Kim SJ, Rah SH, Kang JI, Jung HE, Lee D, Lee HK, Lee JO, Park BS, Yoon TY, Kim HM. Reconstruction of LPS Transfer Cascade Reveals Structural Determinants within LBP, CD14, and TLR4-MD2 for Efficient LPS Recognition and Transfer. Immunity. 2017;46(1):38–50.

40. O’Neill LAJ, Golenbock D, Bowie AG. The history of Toll-like receptors-redefining innate immunity. Nature Reviews Immunology. 2013;13(6):453–460.

41. Gay NJ, Symmons MF, Gangloff M, Bryant CE. Assembly and localization of Tolllike receptor signalling complexes. Nature Reviews Immunology. 2014;14(8):546–558.

42. Han J, Mandal AK, Hiebert LM. Endothelial cell injury by high glucose and heparanase is prevented by insulin, heparin and basic fibroblast growth factor. Cardiovascular Diabetology. 2005;4:1–12.

43. Potter DR, Damiano ER. The hydrodynamically relevant endothelial cell glycocalyx observed in vivo is absent in vitro. Circulation Research. 2008;102(7):770–776.

44. Tsvirkun D, Grichine A, Duperray A, Misbah C, Bureau L. Microvasculature on a chip: Study of the Endothelial Surface Layer and the flow structure of Red Blood Cells. Scientific Reports. 2017;7(October 2016): 1–11.

45. Frey BE a, Miller DS, Jahr G, Sundan A, Espevik IIT, Finlay SBB, Wright SD. of British Columbia, Vancouver, British Columbia, Canada, V6T 1Z3; the *Laboratory of Cellular Physiology and Immunology, The Rockefeller University, New York, New York, 10021; the $Institute for Cancer Research, 7005 Trondheiro, Norway; and the IIInstitu. Response. 1992;176(December).

46. Haziot A, Rong GW, Silver J, Goyert SM. Recombinant soluble CD14 mediates the activation of endothelial cells by lipopolysaccharide. J Immunol. 1993;151(3):1500–1507.

47. Jersmann HPA, Hii CST, Hodge GL, Ferrante A. Synthesis and surface expression of CD14 by human endothelial cells. Infection and Immunity. 2001;69(1):479–485.

48. Lloyd-Jones KL, Kelly MM, Kubes P. Varying importance of soluble and membrane CD14 in endothelial detection of lipopolysaccharide. Journal of immunology (Baltimore, Md.: 1950). 2008;181(2):1446–53.

