## Supplementary Figures for "Heparanase-2 protects from endothelial injury by inhibiting TLR4 signaling"

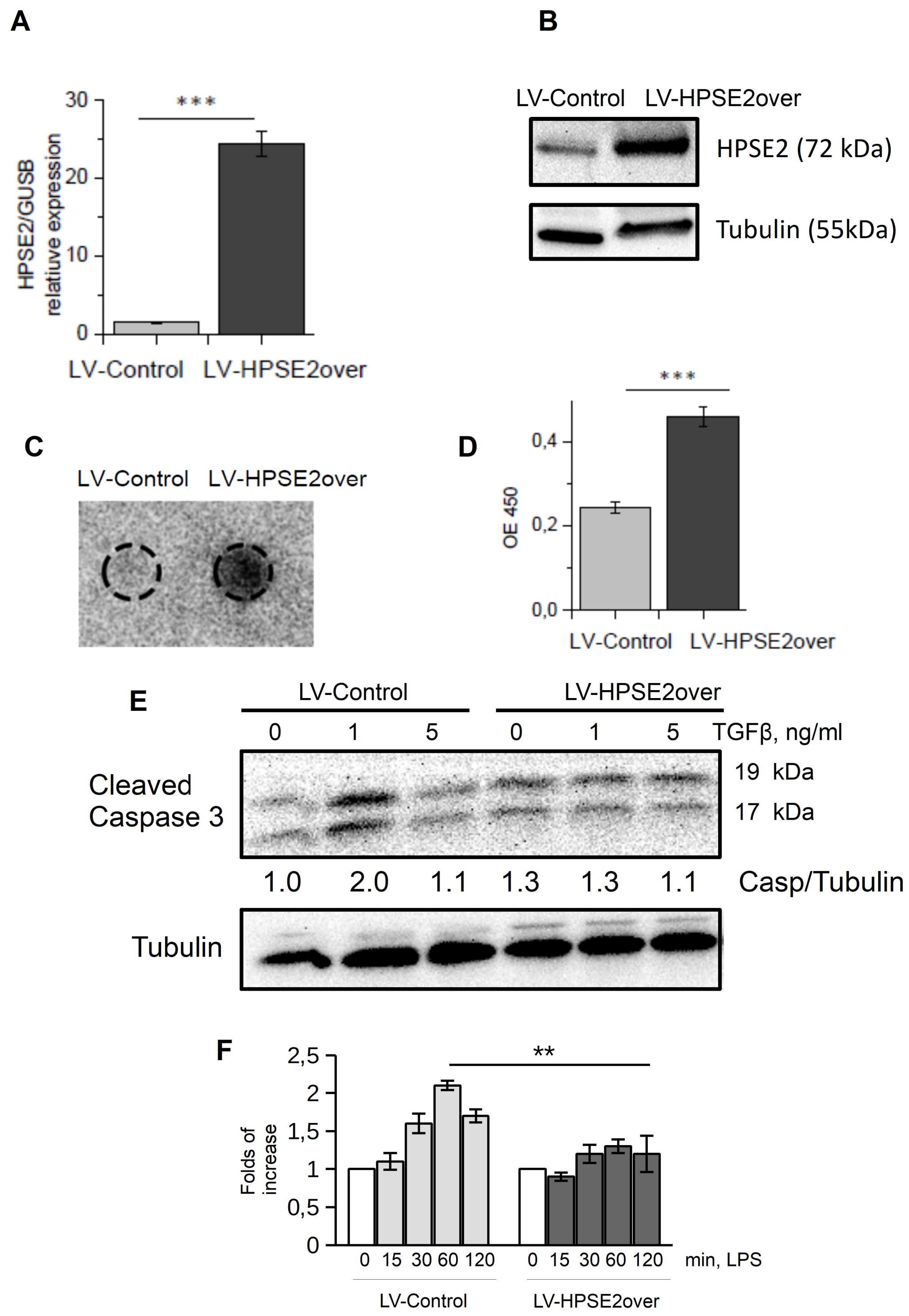


**Supplementary Figure S1. A.** Endothelial cells were infected with control and HPSE2-overexpression lentivirus. HPSE2 expression was analyzed by TaqMan RT-PCR. **B.** HPSE2 expression in lysates of control (LV-control) and HPSE2-overexpressing (LV-HPSE2 over) endothelial cells analysed by western blotting. **C.** HPSE2 presence in conditioned medium of control and HPSE2-over endothelial cells measured by dot blot. **D.** Proliferation of lentivirus-infected endothelial cells was determined by BrdU incorporation assay. **E.** Expression of 17 kDa and 19 kDa fragments of cleaved caspase 3 was analyzed in lentivirus-infected endothelial cells after 16h treatment with TGFβ. **F.** Quantification of western blot experiments as in Fig. 2F.

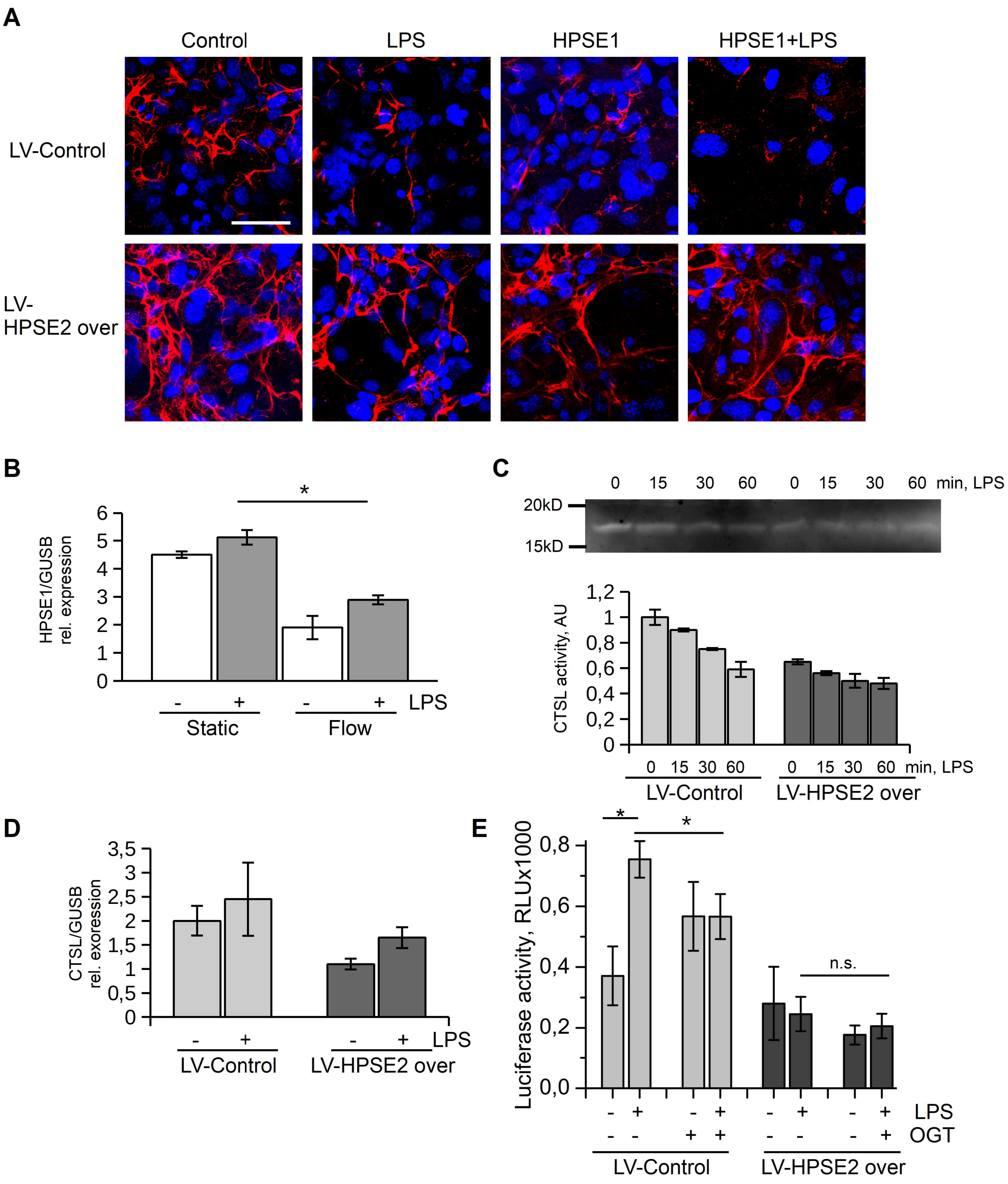


**Supplementary figure S2. Protective effect of HPSE2-overexpression in endothelial cells are mediated by HPSE1 inhibition. A.** Lentivirus-infected endothelial cells were cultivated under flow conditions for 3 days (38.75 dyn/cm$^2$), then treated with HPSE1 for 1 h, and stimulated with 100ng/ml LPS for 3 hrs. Then, cells were fixed and stained with heparan sulfate 10E4 antibody and DAPI. **B.** HPSE1 expression in endothelial cells cultivated in microlfuidic chips under static and flow conditions and stimulated with 100 ng/ml LPS for 3 hrs. **C.** Zymography in gel measurement of intracellular CTSL activity from lysates of endothelial cells. **D.** CTSL mRNA expression was analyzed by TaqMan RT-PCR. **E.** NFκB-driven expression of Gaussia luciferase in lentivirus-infected endothelial cells. Lentivirus-infected cells were pre-treated with HPSE1 inhibitor OGT 2115 and stimulated with 100ng/ml LPS for 1h.

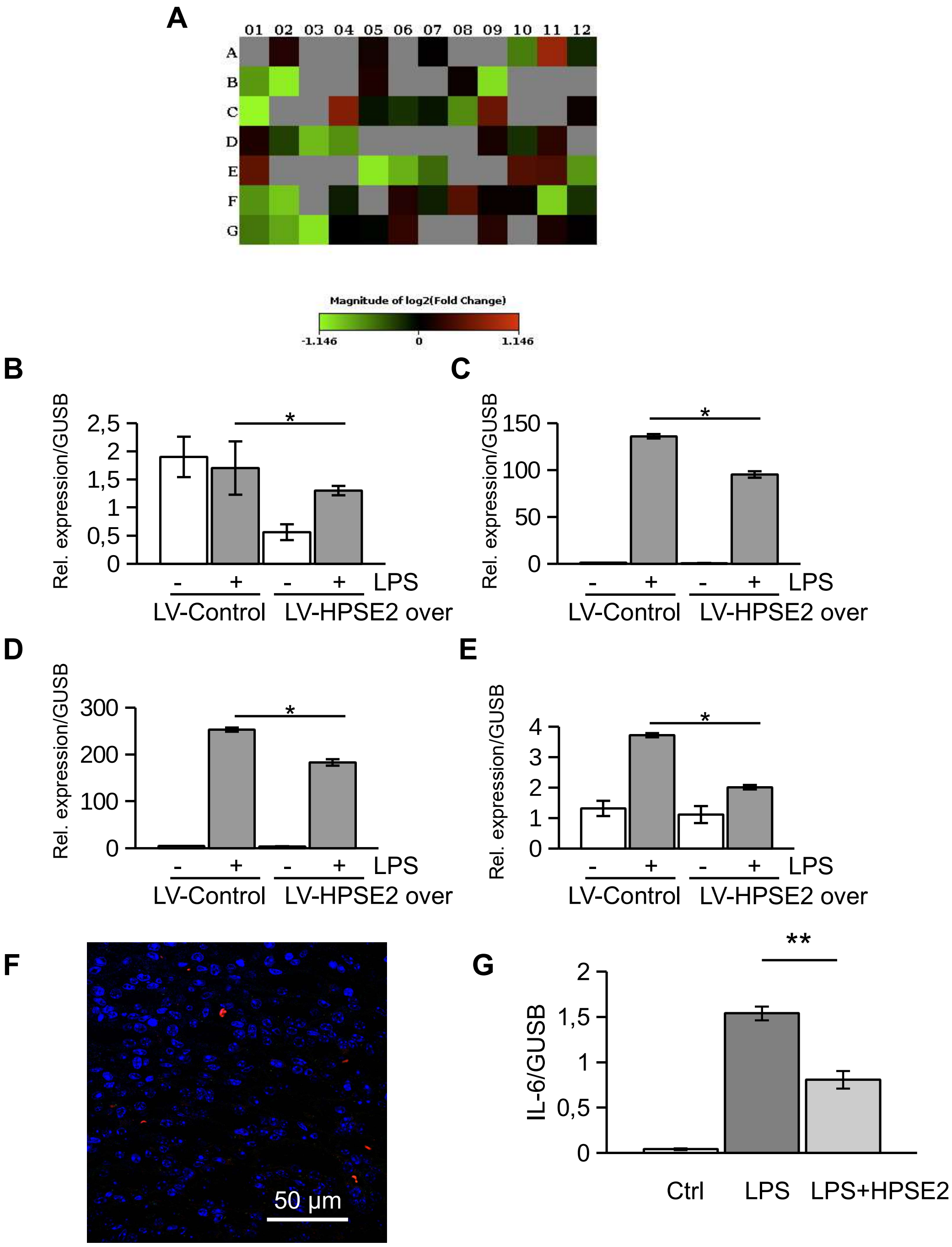


**Supplementary figure S3. LPS-induced response is diminished in HPSE2-overexpressing endothelial cells. A.** Heatmap of the Human Inflammation RT2 Profiler Array (Quiagen) performed from lentivirus-infected and LPS-stimulated endothelial cells. **B-E.** Expression of IFNA1, IFNB1, CSF3, TNFSF10 in the lentivirus-infected and LPS-stimulated endothelial cells was assessed by TaqMan RT-PCR. **F.** IgG negative control of mouse kidney medullary capillaries. **G.** Purified HPSE2 (50µg/ml) was added to endothelial cells in vitro before stimulation with 100ng/ml LPS for 3 hrs. IL-6 expression was assessed by TaqMan RT-PCR.
